# Enhanced Sampling on Domain/Motif Level with Kinetic Accelerated Molecular Dynamics

**DOI:** 10.1101/2025.06.14.659665

**Authors:** Haixin Wei

## Abstract

Molecular dynamics (MD) has become a popular simulation tool in recent years. However, its application is often limited by a timescale problem, due to its femtosecond-level integration timestep. To address this challenge, various enhanced sampling methods have been developed to accelerate system dynamics. Here, we introduce a novel enhanced sampling approach: Kinetically Accelerated Molecular Dynamics (KAMD). By combining the atomic-level accuracy of MD with the diffusive behavior of Brownian dynamics (BD), KAMD significantly improves sampling efficiency on domain/motif level while preserving equilibrium properties. We showed that KAMD is particularly effective in simulating two types of processes: large-scale conformational changes and ligand unbinding events.

## Introduction

Molecular dynamics (MD) is a computer simulation method for calculating the motions of interacting atoms and molecules.^1^ Nowadays, MD simulation has become one of the most widely adopted methods for analyzing and modeling biomolecular structures and interactions.^2,3^ However, classic MD often is limited to short timescales, since its integration timestep is restricted to a few femtoseconds. As a result, it is commonly observed that normal MD is unable to cross substantial energy barriers within a simulation’s lifespan, preventing it from efficiently simulating many complex processes. Thus, a large number of enhanced sampling methods have been proposed over the decades, to accelerate the dynamics of the system and access much longer timescales^4-9^.

The enhanced sampling methods can be broadly classified into two categories. The first category, including umbrella sampling^10^, metadynamics^5^, the weighted ensemble method^11^, etc., utilizes collective variables to represent the degrees of freedom that are of interest. Therefore, the dimensionality of the free energy surface is reduced, and more comprehensive sampling is achieved for these special degrees of freedom. However, the drawback of this type of enhanced sampling method is obvious, that selecting effective collective variables can be very challenging sometimes. It often requires prior knowledge of energy basins and pathways to guide the sampling. As a result, the applicability of such methods is somewhat restrained.

The second category of enhanced sampling methods do not rely on collective variables and comprises a variety of distinct approaches, such as replica exchange dynamics^12^, the mixed Monte Carlo–MD method^4,13^, random accelerated MD methods^14^, etc.^15-18^. These methods try to decrease the free energy barriers or to increase the thermal fluctuation, and therefore accelerate transitions between the different low-energy states^19^. However, reweighting of the resulting trajectories is required to determine configurational probabilities and thermodynamic properties, which can be very challenging.

On the other hand, Brownian dynamics (BD) simulations^20^ allow the simulation of the diffusional motion of molecules in solution, and have been used successfully to explore the kinetics of many biomolecular processes^21-25^. To date, BD has been applied in many biomolecular processes that occur on spatial and time scales out of reach of current MD methods, such as the kinetics of ligand binding to enzymes and receptors,^26-29^ the association and assembly of certain proteins,^30-34^ localization of signaling and target molecules,^35-37^ etc.

However, in BD simulations, the internal structure of the molecules is often treated as rigid, to allow for long timesteps, which is not suitable for many processes that involve flexible biomolecules. For example, intrinsically disordered proteins (IDP), a type of protein that lacks a fixed or ordered three-dimensional structure, sample a large area of conformation space, and cannot be treated as rigid.^38-45^ Another evident example is the ligand binding on a flexible target, in which conformational change will be induced or adopted in the receptors.^46^ This type of processes happens in variety of systems, including G protein coupled receptors,^47-49^ kinases,^50-52^ ion channels,^53,54^ viral proteins,^55-58^ etc. As such, the need of a new generation of BD simulation programs which can realize both long simulation time and molecular flexibility is prominent.

In this work, we will combine the atomic accuracy of MD and diffusion behavior of BD to give a brand-new enhanced sampling method, Kinetic Accelerated Molecular Dynamic (KAMD) simulation. Comparing to the first category of enhanced sampling methods discussed above, KAMD does not utilize collective variables, allowing convenient application to many more systems. Besides, unlike high temperature or lower potential energy, KAMD utilizes special velocity distributions in classic MD to accelerate kinetics. The velocity distributions represent events that, although rare, do occur in standard MD simulations, allowing KAMD to faithfully reproduce MD configurational sampling over long-time scales. We will show that KAMD is particularly good at simulating two types of structural events, large-scale conformational changes and ligand unbinding events. With full development, KAMD should prove widely useful for scientific discovery in molecular biology.

## Method

### 1. Kinetic accelerated molecular dynamic simulation procedure

Kinetic Accelerated Molecular Dynamic (KAMD) simulation is designed to adapt features of Brownian Dynamic simulation into classic molecular dynamic simulation, to accelerate sampling of selected processes.

In a BD system, several bodies interact and move, to simulate conformation changes, binding/unbinding events, etc. BD simulation is commonly used for biomolecular systems since many proteins/nucleic acids can be safely approximated as assemblages of independent bodies. By making a rigid-body approximation, BD simulation can ignore atomic details and can simulate longer time scales compared to MD simulation. Therefore, KAMD will mimic the BD behavior and boost the domain/motif level of configurational sampling of classical MD systems.

KAMD requires the user input of “region of collective motion” (denoted as “region” for following part, up to six for current version of the program), which are some parts of the molecule simulated. These “regions” resemble the rigid bodies in BD systems, and their intramolecular motions are not of primary interest. In biomolecular systems, these “regions” widely exist and are easily recognized, such as a domain, a subunit, or a motif, etc. The basic idea of KAMD is to add one layer of BD motion to these “regions”, while on the atomic level, keeping the normal MD simulation.

The BD movement of these input “regions” will be added by the following procedure:

1. For each “region”, its center of mass velocity *ν*_*com*_ will be tracked in the simulation. The tracking happens according to an input quantity, BD step; BD step controls the acceleration level of KAMD, and the highest acceleration is related to a BD step value of 500fs, as this is about the correlation time of a water molecule. For slower acceleration, BD step value can be turned up, for example, to 1ps. *ν*_*com*_ will be marked as direction-changed if the dot product between this BD step and last BD step is less than zero:

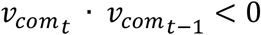 Once all the input “regions” have changed direction twice, the system motion will be deemed to be oscillating, and the system has entered normal MD status. At this point, we assume the system no longer does any meaningful “region” level of conformational sampling and waits for the next acceleration cycle.
2. Starting a new acceleration cycle by randomizing the velocities of the entire system.
3. After randomizing all velocities in the system, the program will find and mark two types of atoms: 1. solvent molecules, including free ions and waters, within 5 Å (first solvation shell) of each input “region” and 2. surface atoms of each “region” that are within 3 Å of any solvent. The first type of atom, in the surrounding solvent molecules, will be given an ownership to the closest “region”. The second type of atom, of course, belongs to its own “region”.
4. For each of the “regions”, one random direction 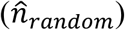 will be generated. The two types of marked atoms that belong to this “region” will have their velocities reoriented, based on 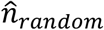, by the following relation:

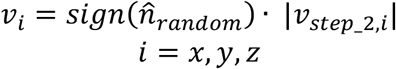

By doing this step, “regions” gain a BD motion since all the marked velocities of each “region” are approximately oriented to the one chosen direction, 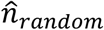.

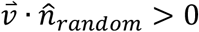

Noted that the velocity magnitude distribution is preserved:

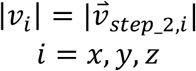

A schematic illustration of step 4 is shown below in figure 1.

**Figure 1.**
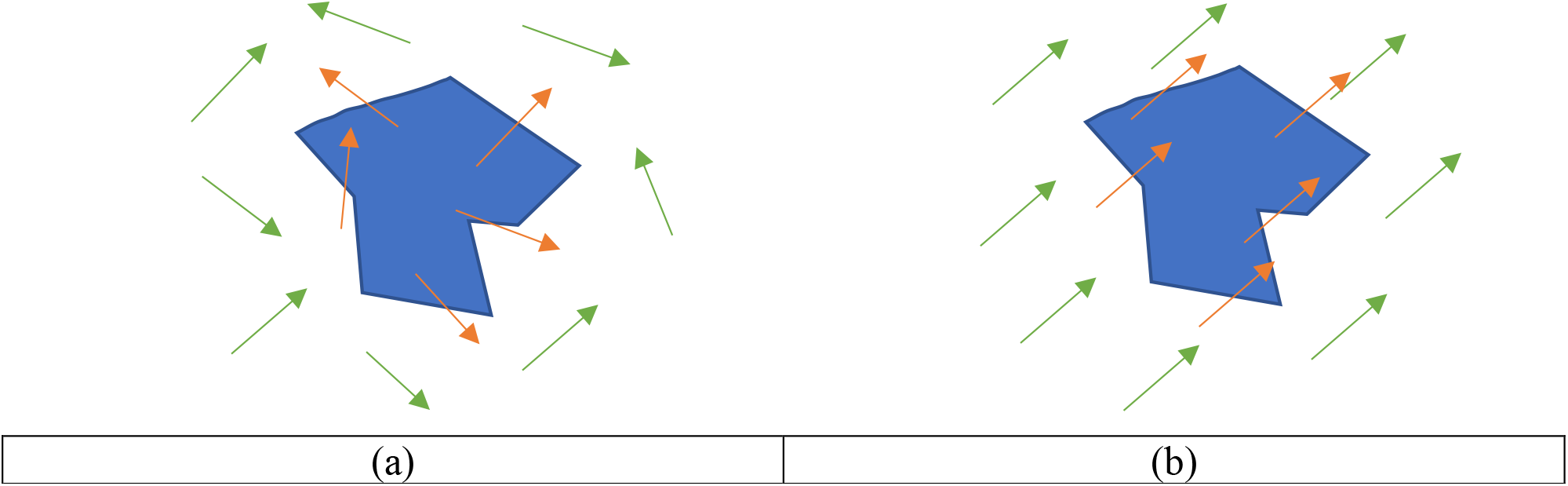
The reorientation of velocities in the step 4. (a) The velocity distribution of marked atoms before step 4. (b) The velocity distribution of marked atoms after step 4. The blue shape represents an input “region”, the green arrows represent velocities of first type of marked atoms, and the red arrows represent velocities of second type of marked atoms. Step 4 is the key step for KAMD acceleration. As we showed above, KAMD utilizes a rare velocity distribution (aligned to one direction for each “region”) to accelerate changes in the system. Although this special type of velocity distribution is a rare event in normal MD, it will exist if a MD simulation lasts long enough. Thus, KAMD trajectories are authentic, as they are just reproducing long-time behavior of classic MDs. This is a major advantage comparing with other enhanced sampling methods.
5. Resume the simulation with the new velocity distribution and monitor the velocity correlations.

### 2. KAMD preserves conformational free energy

In the above procedure, the “region” level of movement of the simulated system will be accelerated, but the conformational free energy of the macromolecules will be preserved.

Free energy is the most important property of a simulation, as most of the measurables are related to it. We provide a theoretical proof in **Supporting Information** and an Alanine dipeptide test in **Results and Discussion** section. In a word, it is safe to use KAMD for quantitative analysis and one can compare the results with experiments directly.

### 3. KAMD preserves time evolution approximately

In usual biomolecular simulations, people sometimes are interested in studying transition pathways, which is related to the system time evolution in conformational space. We can prove, by both theory and examples, that KAMD is able to preserve the system time evolution, in a short period of time. The detailed proof is provided in **Supporting Information**, and example is provided in **Results and Discussion** section.

## Results and Discussion

KAMD is a general-purpose molecular simulation tool. Based on current development, KAMD is mostly advantageous in simulating two type of processes, conformational sampling and ligand unbinding. In the following sections, we will use examples to illustrate the performance of KAMD. First, we will use an Alanine dipeptide test case to validate some basic physical properties of KAMD. Then, we will present three examples: TSPI polypeptide, thrombin, and IgE-Fc, to show the accelerated sampling capability of KAMD. Lastly, we will use 12 HIV-1 protease inhibitor systems to show how KAMD can benefit ligand unbinding studies.

### 1. Alanine dipeptide

Due to its simplicity, Alanine dipeptide is a popular model for studying the behavior of a simulation tool. Here, we will use the Alanine dipeptide test case as an example to show three key features of KAMD: 1. KAMD preserves the free energy surface 2. KAMD samples faster than MD 3. KAMD has similar time evolution as MD, in a short period of time.

The simulation details that coincide with MD are provided in **Supporting Information**. Here, we provide the KAMD specific inputs, which are the “regions of collective motion”. We show in Figure 2 (a) the two input “regions” used: left eight atoms (left rectangle) and right eight atoms (right rectangle). The choice of the input “regions” should be obvious, as the conformation of Alanine dipeptide is primarily determined by the relative motion of these two “regions”. The BD step value was set to 500fs.

**Figure 2.**
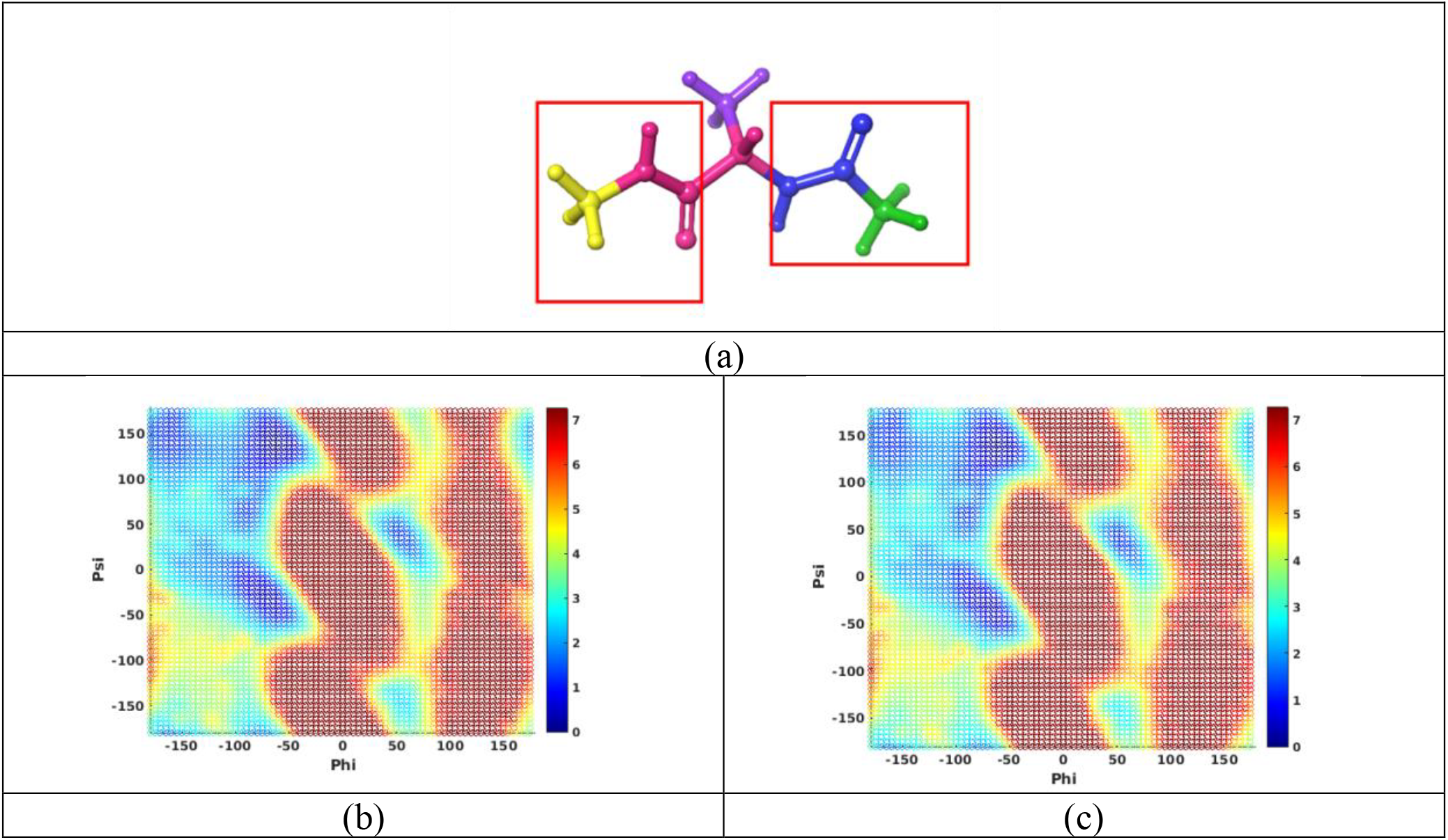
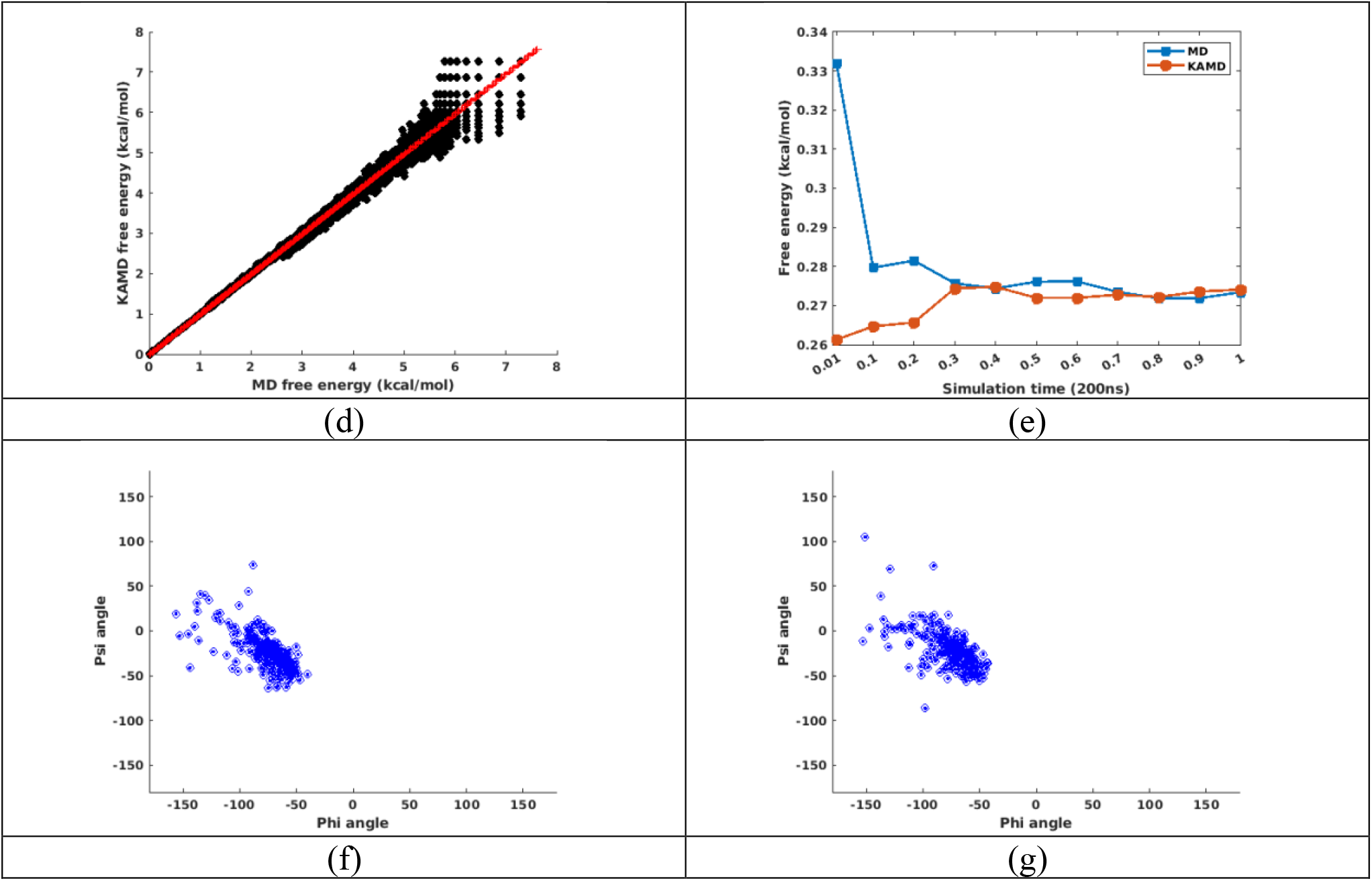
Test results of Alanine dipeptide. (a) Alanine dipeptide structure, red rectangles indicate two input “regions”. (b) Ramachandran plot of MD simulation (c) Ramachandran plot of KAMD simulation (d) Free energy fitting plot of entire free energy surface (e) Free energy convergence plot for *ε* conformation: Phi = -70° and Psi = 130°. (f) Conformation distribution after 2ps MD simulation, from *α* conformation: Phi = -75° and Psi = -20°. (g) Conformation distribution after 2ps KAMD simulation, from *α* conformation.

To demonstrate that KAMD preserves free energy, we first showed the Ramachandran plots of Alanine dipeptide from both MD and KAMD simulation. In figure 2 (b) and (c), the two Ramachandran plot are extremely similar. Both the number and position of the “hills” and “valleys” are same. Both of the plots show two major favorable conformation: the most favorable conformation, the *ε* conformation, around Phi = -70° and Psi = 130°, and the second favorable conformation, the *α* conformation, around Phi = -75° and Psi = -20°. Both of the plots showed the less favorable conformation around Phi = -150° and Psi = -130°, which is the *β* conformation.

We also show the correlation plot for all the data points on the Ramachandran plots in Figure 2 (d). The slope of the curve is 0.995, showing excellent correlation. The errors are only present in high free energy area, around 6 to 8 kcal/mol, which is not sufficiently sampled by MD (and probably KAMD, too).

To prove that KAMD samples the conformational space faster than MD, we plot the free energy convergence plots, as shown in Figure 2 (e) and Figure S1. Clearly, from the figures, we can see that the free energy of KAMD and MD converge to the same value as simulation time goes by. This is also a proof of the free energy conservation property. However, when we look at shorter time scale, for example, the first data point, 2ns, in Figure 2 (e), we can see clearly that MD value is much more deviated from the convergence value. The error of the free energy of 2ns KAMD simulation is about the same as the error of the 20ns MD value, showing that KAMD is about 10 times faster than MD in Alanine dipeptide situation.

We want to mention an important point here. The acceleration of KAMD is dependent on the input “regions” of the system. Here, for the Alanine dipeptide simulation, the “regions” only contain 8 atoms each, so it displays a comparatively slow acceleration, about 10 times. We will show later, as the simulation systems get larger, the KAMD can accelerate by thousands and even more fold.

Lastly, we show the time evolution plot in Figure 2 (f) (g) and Figure S2. Time evolution, as discussed before, can be of interest in some transit process, such as conformational change pathway. In Figure 2 (f) (g), we show the conformational distribution after 2ps simulation, starting from the *α* conformation (Phi = -75° and Psi = -20°). The distribution is very similar for KAMD and MD, with a slightly larger area covered by KAMD simulation. The plot for 40ps, 100ps and 200ps are provided in Figure S2, and they all showed similar behavior. This proved that KAMD can also preserve time evolution on a short time scale. At 200ps, the distribution is close to Ramachandran plot, showing approaching equilibrium. Thus, the distribution of longer time is not relevant.

### 2. TSPI polypeptide

Proline related cis-trans transformation, as shown in Figure 3 (a), regulates many key interactions in molecular biology, and here we used a TSPI polypeptide to study this conformational change. It should be noted that this process happens very slowly, at millisecond scale at 400K^59,60^ and way beyond the capability of classic MD.

**Figure 3.**
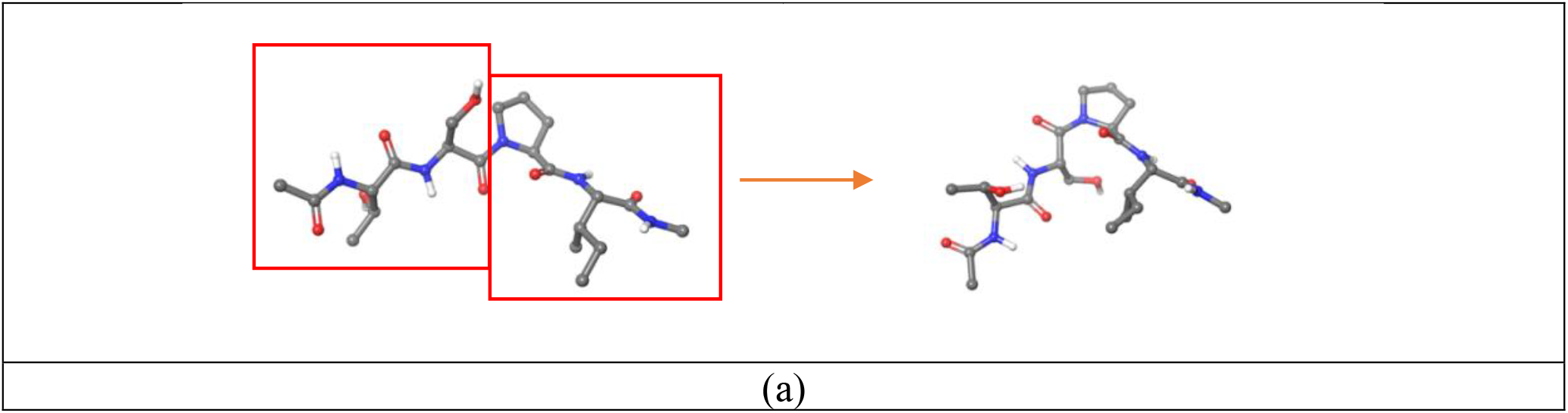
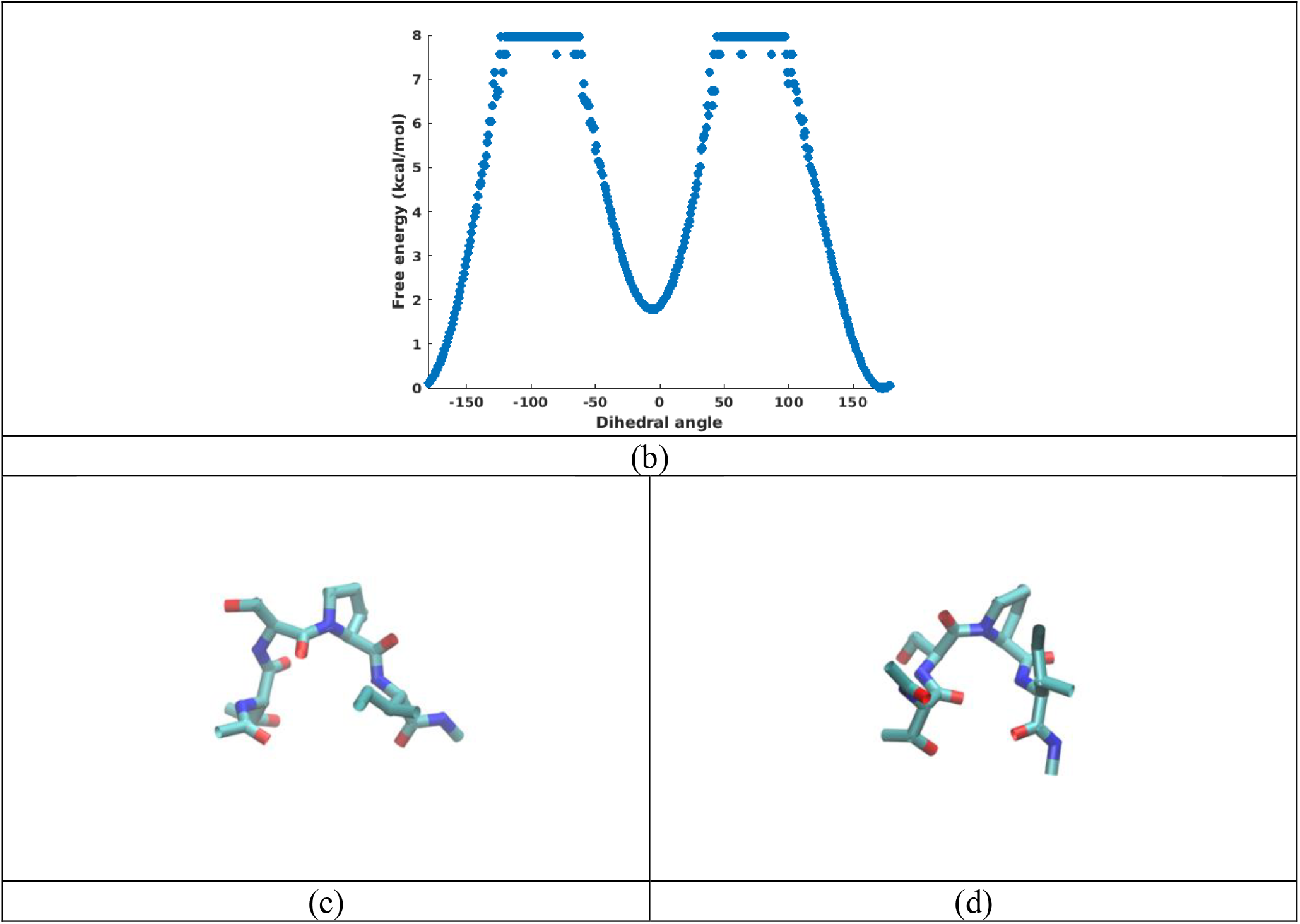
KAMD simulation results of polypeptide TSPI. (a) Schematic illustration of trans-to-cis conformational change. The red rectangles indicate two KAMD input “regions”. (b) Free energy plot of dihedral angle in Ser-Proline peptide bond. (c) The last simulation frame before the trans-cis conformational change happened. (d) The first simulation frame after the trans-cis conformational change happened.

There are 2 input “regions” for TSPI system, as shown in Figure 3 (a) in red rectangles: the first two residues, TS, and the last two residues, PI. The choice is decided based on the process we are interested in. For the cis-trans transformation TSPI, the above choice should be obvious and optimal. The BD step value was set to 500fs. Other MD simulation details are provided in **Supporting Information**.

The most important test is, of course, whether the cis-trans transformation happened. In Figure 3 (b), we plotted the conformational free energy of the Serine-Proline peptide bond dihedral angle. We can see that, the transformation happened and a second free energy valley appeared, around 0 dihedral angle. The transformation only happened in one of ten independent trajectories, at around 1.1μ*s*. After the transition happened, the polypeptide stayed at cis conformation stably until 2μ*s*.

The acceleration of TSPI cis-trans conformational change cannot be obtained by comparing KAMD results with MD results, since the conformational change never happened in MD simulation. But we can compare the simulation results with similar experiment results, which is millisecond phenomenon. Thus, we can say that the acceleration of KAMD is about hundreds to thousands fold, for TSPI cis-trans conformational change situation.

We also show the structural details right before and after (2ps away from each other) the conformational change, in Figure 3 (c) (d). From the two structures, we can infer how the conformational change happened. Imagine that you are clamping the two ends of the peptide, Threonine and Isoleucine, with your two figures. At conformation Figure 3 (c), the “clamping” will force the two ends to get closer. However, the peptide will want to resist this “clamping” and a tension will be created and concentrate to the middle of the peptide, which is exactly the S-P peptide bond. If this tension is strong enough to cross the transition barrier, the conformational change will happen and the structure Figure 3 (d) forms. This transition mechanism is supported by Figure S3, which measures the distance between the two C-*α* atoms of Threonine and Isoleucine. From Figure S3 (b), we can clearly see that the two C-*α* atoms come to their local closest distance, right before the conformational change happens, and after the transition, the distance stays at a low value for a little longer.

In **Supporting Information** and the Alanine dipeptide test, we have shown that the transition pathway of KAMD is approximately preserved. As can be seen from Figure S3 (b), the dihedral transition happened within 10 ps of the distance dropping in the trans-conformation. Thus, the geometric character of the transition (i.e., the reaction pathway) is roughly preserved in KAMD, similar to what will be seen in a long enough MD simulation.

### 3. Thrombin

Thrombin is a serine protease enzyme that plays a crucial role in blood clotting. There is a known “open-close” conformational change related to its activation mechanism. When thrombin is free in solution, there are two major conformations. In the inactive conformation, the side chain of W215 and backbone of 215-219 collapse in the active site, forming the “closed” conformation. On the other hand, in the active conformation, the W215 indole ring and the entire 215-219 segment move aside, allowing the active site to be wide open, forming the “open conformation”. The rate of this conformational change is estimated to be about 400/s, which is beyond the capability of classic MD simulation. Here, we show that KAMD can reproduce this conformational change.

The input “regions” for thrombin are two: N-terminal domain and C-terminal domain, as we show in Figure 4 (a). The two domains are easily recognized in the structure and require no pre-knowledge of the “open-closed” conformational change. The starting structure was taken from PDB ID 3BEI, which is a “closed” conformation. Our purpose is to see if KAMD can transform this starting structure to the “open” conformation, which is represented by structure of PDB ID 1SG8. The BD step value was set to 1ps. Other MD simulation details are provided in **Supporting Information**.

**Figure 4.**
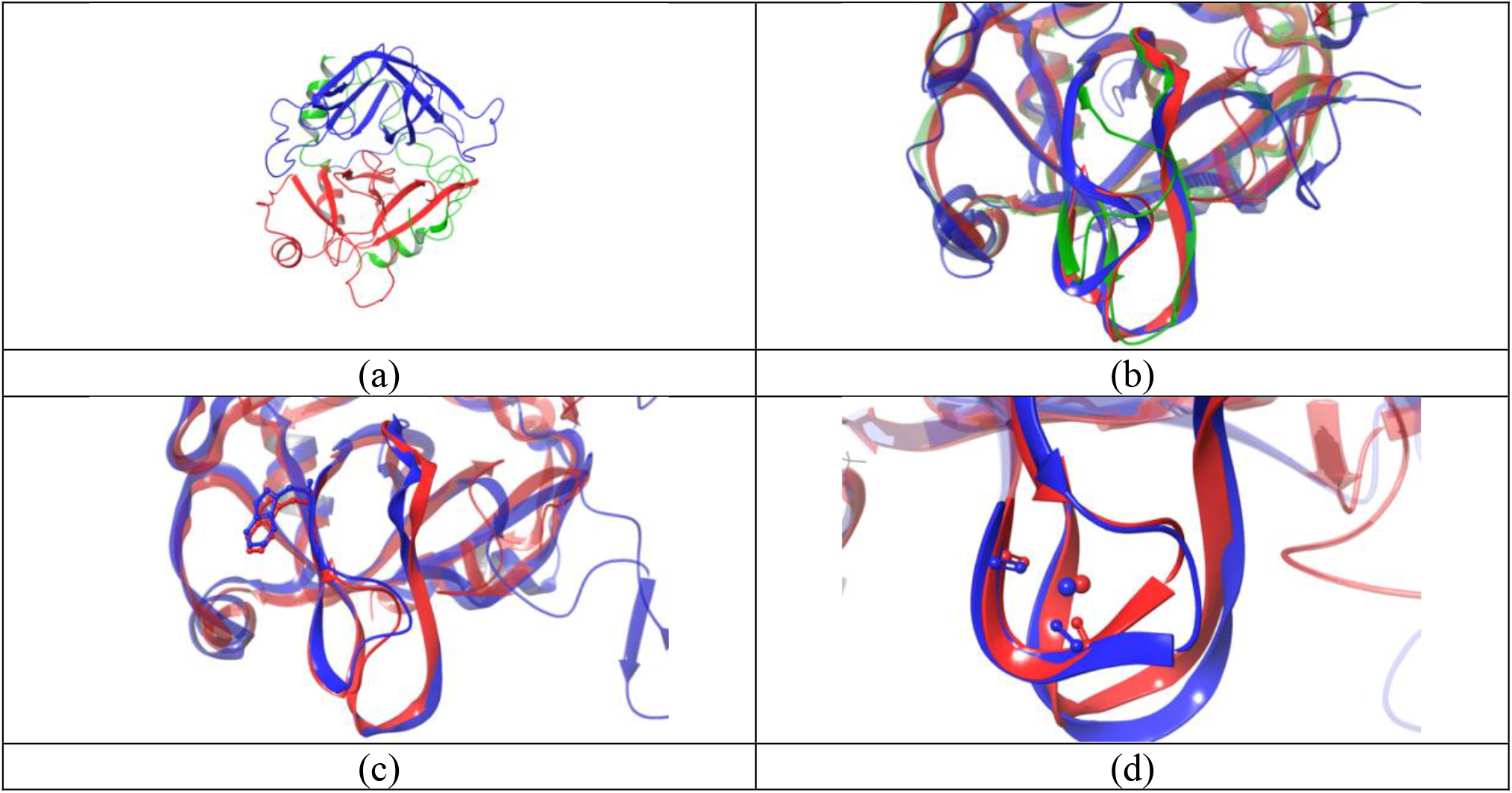
Simulation results for protein thrombin. (a) Structure of thrombin. Blue and red color indicate two input “regions” of KAMD. The green part is not in any input “region”. (b) Active site structure comparison between starting structure, (green, PDB ID 3BEI), target structure (red, PDB ID 1SG8), and the simulated structure that showed smallest active site RMSD (blue). (c) The smallest RMSD structure when the W215 site fully opened (blue), and the target structure (red). (d) The smallest RMSD structure when sodium ion is bound at crystallized position (blue), and the target structure (red).

The result showed clearly that KAMD is able to reproduce this millisecond timescale phenomenon. Figure 4 (b) showed the frame from KAMD trajectories that is of the smallest active site RMSD value. The RMSD value, comparing with 1SG8 structure, is 0.995 Å, which is smaller than most of crystallization accuracy. This is suggesting that the simulated structure is of a similar quality as a crystalized structure.

There are two important interactions in 1SG8 that mark the “open” state of the protein. The first one is the hydrogen bonding between W215 and M180. This hydrogen bonding is a sign of fully opened W215 indole ring. As we can see from Figure 3 (c), this interaction is realized in the KAMD trajectories. The W215 side chain has moved to its crystalized position, and the active site is completely exposed. The second important interaction is that some sodium ions may bind at a pocket below the active site, as we showed in figure 4 (d). In the starting structure, 3BEI, there is no sodium ion at that position. However, since there are free sodium ions in the solution, there is possibility that some sodium ions will find this binding site during the simulation. Here, we showed in Figure 4 (d), there are indeed sodium ion binding at that place. The distance between the ions in the best fitted structure and the crystal structure is about 0.5 Å. Again, this behavior clearly proves that KAMD can faithfully reproduce the thrombin “open-close” conformational change.

### 4. IgE-Fc and IgE-Fc-sFc*ε*RI*α* complex

Immunoglobulin E (IgE) binding to its high affinity receptor, FcεRI, through its Fc region (IgE-Fc). The high affinity binding is a key step in allergic diseases, and its half dissociation time is about 14 days.^61,62^ X-ray crystallography study of both free IgE-Fc and IgE-Fc-sFcεRIα complex showed a bent structure.^63^ However, their behavior in solution is not clear. There have been fluorescence studies on the subject, showing that bending exists in solution also.^64^ Here, we use KAMD to study the system.

The KAMD simulation was carried on two systems: free IgE-Fc and IgE-Fc-sFcεRIα complex. The two simulations will be used to compare with the fluorescence experiments. A classic MD simulation was also done for free IgE-Fc system, to show that MD is not sufficient for this problem.

The input “regions” for two KAMD systems are the same, which is six. The IgE-Fc is a homodimer formed by six subdomains, so naturally, the six domains are the six input regions. The sFcεRIα protein was filled with both N- and C-terminal tail, using AlphaFold database, and buried into a membrane. The starting structures are taken from PDB ID 2WQR and 2Y7Q, respectively, for free IgE-Fc system and IgE-Fc-sFcεRIα complex system. The BD step value was set to 1ps. Other simulation details are in **Supporting Information**.

First, we compared normal MD trajectories and KAMD trajectories of free IgE-Fc system, for conformational sampling. We show the principal component analysis (PCA) results in Figure 5 (b) and (c). It can be clearly seen that KAMD samples a much larger area than MD. By simply counting the pixels, the area covered by KAMD trajectories is about 5 times of that of the MD trajectories. Also, when we cluster the trajectories of the MD simulation and generate one single representative structure, as we show in Figure S4, it can be seen that the representative structure is very similar to the crystal structure, with a RMSD value of only 1.9 Å. This means that the MD simulations never really leave the crystalized conformation and never approach the solution conformations. Thus, simply using conventional MD results to compare with fluorescence experiments is not helpful, because there is not much more information than in the crystal structure.

**Figure 5.**
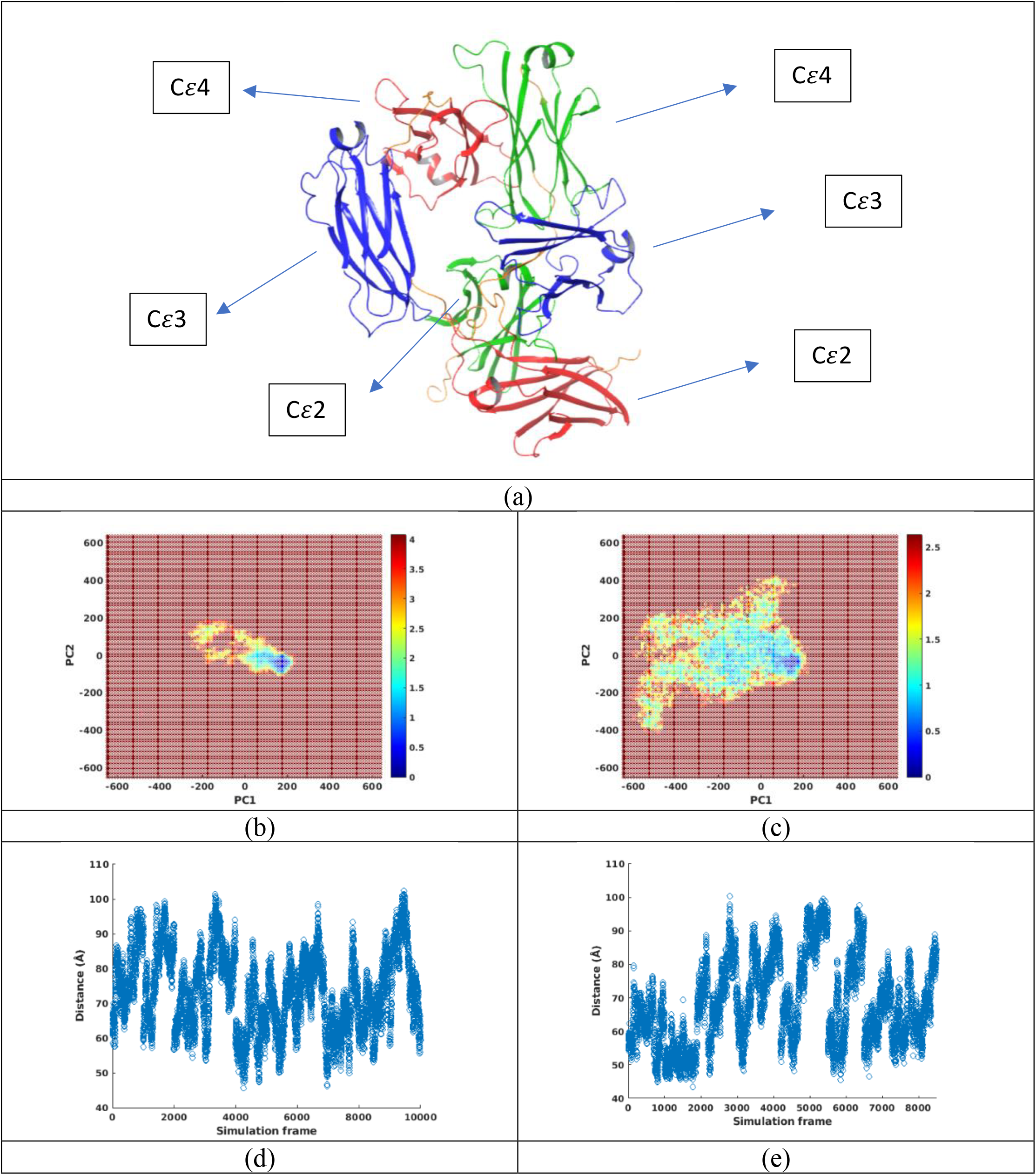
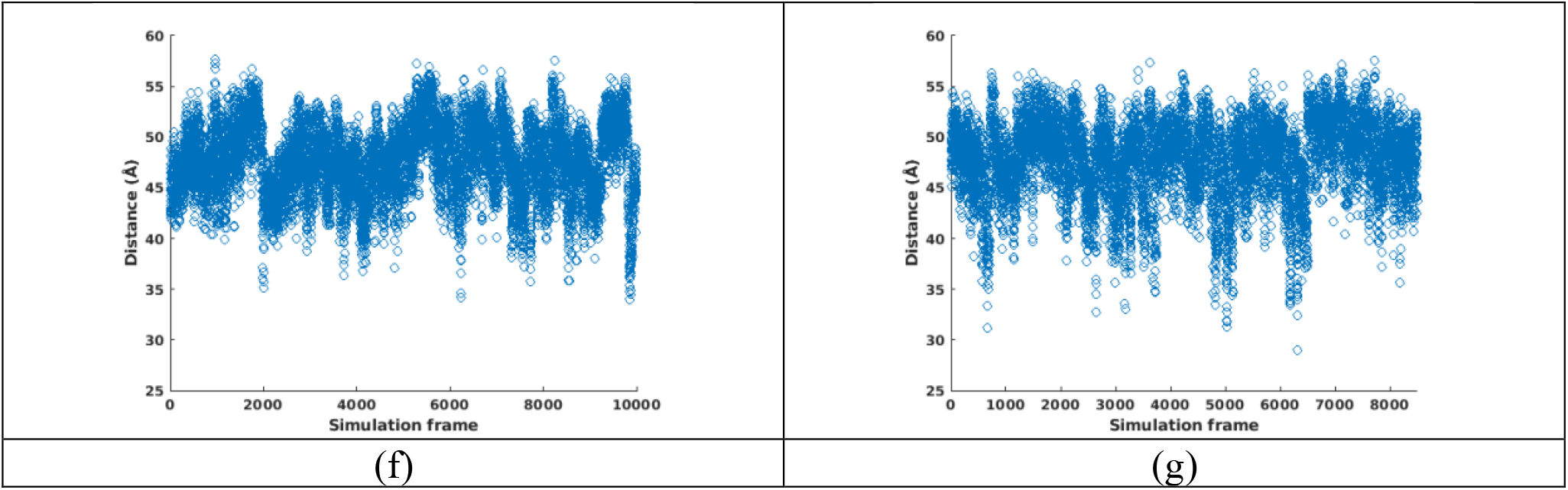
Simulation results of free IgE-Fc and IgE-Fc-sFcεRIα complex systems. (a) Input “regions” for the two systems, using free IgE-Fc system as an example. Each red, blue, and green region represents one domain of IgE-Fc and is used as an input “region” in KAMD. The yellow parts are linkers and are not in any input “region”. (b) PCA result of MD of free IgE-Fc system. (c) PCA result of KAMD of free IgE-Fc system. (d) Distance between Ser542 and Phe229 of the free IgE-Fc system. (e) Distance between Ser542 and Phe229 of the IgE-Fc-sFcεRIα complex system. (f) Distance between Phe229 of A/B chain of the free IgE-Fc system. (g) Distance between Phe229 of A/B chain of the IgE-Fc-sFcεRIα complex system.

On the other hand, KAMD trajectories have really pushed the systems into solution state behavior. The most populated structure from clustering the KAMD trajectories is of a RMSD value of 12 Å (Figure S4), showing that the KAMD has transformed the system conformation far away from crystal structure. Actually, even further, KAMD has reached simulation limits for some trajectories. In 10 KAMD trajectories of IgE-Fc-sFcεRIα complex, 3 showed complex dissociation and 1 showed C*ε*4 dimer uncoupling (Table S1). These major structural events mark critical points of conformation sampling and should be end points for some KAMD trajectories. For example, for later comparison between free IgE-Fc and IgE-Fc-sFcεRIα complex KAMD simulations, we included the frames that C*ε*4 dimer uncoupled but not those complex-dissociated ones. This is because C*ε*4 dimer uncoupling is not forbidden in reality, when IgE-Fc-sFcεRIα complex exists. Thus, all frames that are before complex dissociation should be included when consider the conformations of IgE-Fc-sFcεRIα complex in solution. The detailed information about frame selection can be found in Table S1.

To reach some major structural events, like complex dissociation here, is critical for statistical comparison with experiments. This is because only when you have reached the simulation limits, you can know that your simulation is long enough for finishing the sampling and converging the ensemble statistical quantities. This is the only way that you know your data is not biased towards conformations near the starting structure. Thus, using most of MD trajectories to compare with experiments is not appropriate, because the ensemble statistics is far from converged. Even if the data appear to be internally converged, it is likely that the convergence is around a local minimum rather than global, so still cannot truly reflect the experimental conditions. Here, we showed that KAMD can reach the simulation boundaries and therefore prove sufficient sampling. Then, the results can be safely employed to compare with experiments.

Knowing that KAMD reflects the true protein conformations under solution conditions, we can now compare the results with fluorescence experiments (note that 8500 frames were selected before complex dissociation happens, see Table S1). There are two major conclusions from the experiments.^64^ First, the bending of the IgE-Fc protein increased when forming a complex with sFcεRIα. To confirm this behavior in KAMD simulation, we calculated the distance between the C-terminus Ser542 and N-terminus Phe229, as we plotted in Figure 5 (d) and (e). The average values are 73.5 Å and 68.4 Å, for free IgE-Fc system and the complex system respectively. This means the bending is indeed increased for the IgE-Fc-sFcεRIα complex system and it is consistent with the fluorescence experiments. The 5.1 Å distance difference also agrees with the fluorescence experiments, because the 6.9% distance difference of the two systems agrees with the 7.3% fluorescent intensity difference in their calculation^63^.

The second major conclusion of the fluorescence experiments is that the C*ε*2 dimer does not dissociate in solution, when binding with sFcεRIα. This is also confirmed in KAMD simulation, as we showed in Figure 5 (f) and (g) the distance plot between two N-terminus Phe229 residues. The distributions in the plots are very similar and the averaged values are 47.7 Å and 47.8 Å, for free IgE-Fc system and the complex system respectively. The difference is only 0.1 Å and can be treated as negligible, agreeing with the fluorescence experimental result. However, C*ε*4 dimer showed uncoupling behavior in the IgE-Fc-sFcεRIα complex system but not in the free IgE-Fc system, indicating different free energy landscapes for the C*ε*4 dimer in the two systems.

### 5. HIV-1 protease inhibitors

HIV-1 protease is a popular drug target and has been widely studied in recent decades. There are 12 inhibitors in the literature for which both crystal structures and k_off_ rates are available, making it a perfect system for testing KAMD ligand unbinding simulations. Here we show the simulation study below.

There are two types in the 12 inhibitors. The type-1 inhibitors, including three structures, interact with the protease directly. The detailed active site view can be found in Figure S5 (b). Differing from type-1 inhibitors, the other nine type-2 inhibitors have an important water-mediated interaction with Ile50 in the binding site, as shown in Figure S5 (d). Because in KAMD, the system movements are accelerated, and different systems will have different acceleration, time-related comparisons can only happen relatively between similar systems. So, the KAMD simulation was also divided into two groups, to compare their simulated dissociation times.

The input “regions” are three, for both types of inhibitors: subunit A/B and the inhibitor itself (Figure S5 (a) and (b)). However, the type 2 inhibitor system requires one more layer of complexity: the water that mediates interaction with Ile50 must be treated as a special water molecule. The reason is that in KAMD, atoms in the input “regions” that are near the solvent will get their velocities reoriented (see **Method**). However, from the crystal structure, we know that this special water molecule is an interacting water and should not be given the capability to reorient nearby atom velocities. Thus, in type-2 inhibitor systems, this water was treated as a second ligand to differ from normal solvent waters. Also, a distance restraint was applied to keep the hydrogen bonding between this water and the inhibitors. This restraint is to prevent water exchange happening between this special one and other normal solvent waters. Obviously, if other water takes this place instead, the velocity reorientation problem happens again. So, the restraint is necessary, and without it, all the type-2 inhibitors quickly dissociated. On the other hand, the restraint should have very little effect on KAMD dissociation simulation, as the restraint energy is non-zero for less than 1/1000 frames.

The BD step value was set to 500fs, and all other simulation details are in **Supporting Information**.

We list the KAMD averaged relative dissociation time of the two types of inhibitors in Tables 1 and 2. The dissociation of the inhibitor was determined by ligand RMSD value for each trajectory. Once the RMSD value reaches 20 Å, the inhibitor was deemed to be dissociated, and that frame time was recorded as the dissociation time for the trajectory. Then, we take all the dissociation times and average them to give the final dissociation time results, which are reported in Tables 1 and 2. Note that those trajectories in which inhibitor dissociation did not happen are not included in the average, so strictly speaking, we are calculating the lower bound of dissociation time. The value “>1000” in the Tables 1 and 2 means that, in all the 10 trajectories, not even one dissociation happened, so there is no estimation of the dissociation time. The ligand RMSD plot can be found in Figure S6.

**Table 1.** KAMD averaged relative dissociation time of type-1 inhibitors.

| PDB ID | $k_{\text{off}}$ rate/s | KAMD dissociation time (ns) |
| --- | --- | --- |
| 1qbs | 83.3 | 666 |
| 1g2k | 0.0697 | > 1000 |
| 1g35 | 0.0685 | > 1000 |

**Table 2.**
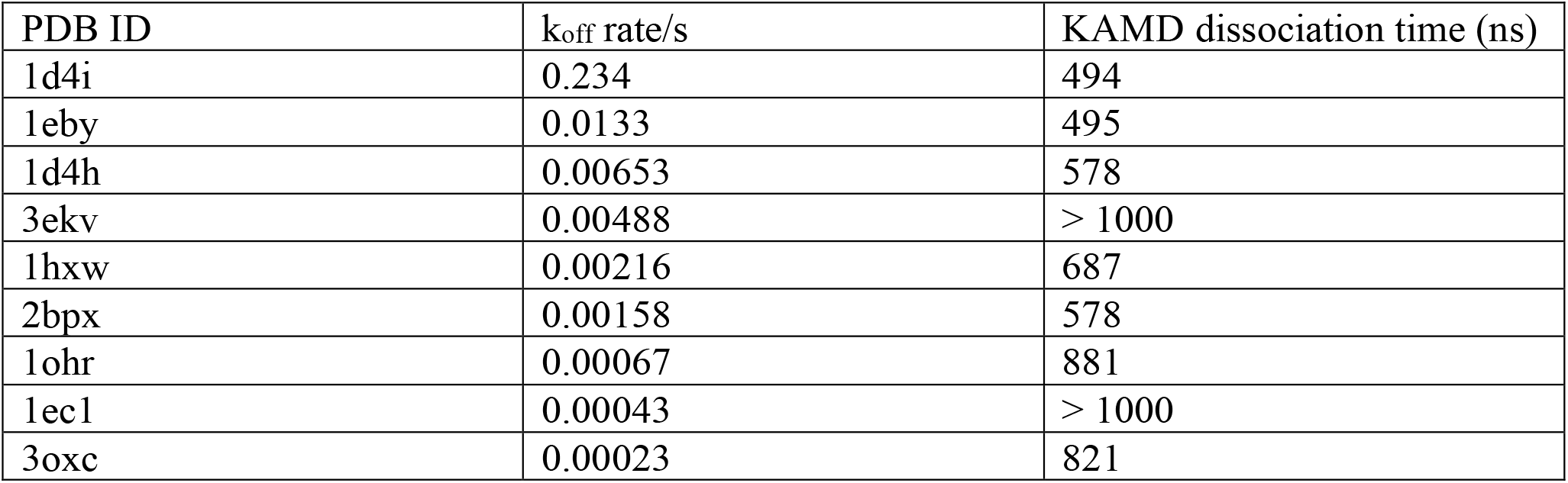
KAMD averaged relative dissociation times of type-2 inhibitors.

It is clear, from the results in Tables 1 and 2, that KAMD is capable of reproducing the inhibitor unbinding process. For type-1 inhibitors, 1qbs showed dissociation because it is of a very high k_off_ rate. The other two, 1g2k and 1g35, showed almost the same behavior for both k_off_ rate and dissociation time, proving the correctness of KAMD’s prediction. Things are bit more complicated with type-2 inhibitors, but we can clearly see the trend, that KAMD dissociation time increases as k_off_ drops. There are two inhibitors that do not show dissociation in the simulation, 3ekv and 1ec1, and it seems 3ekv is an outlier. The reason for this behavior is probably the inhibitor parametrization. We reported the CHARMM ligand parametrization penalty values in Table S2, and 3ekv and 1ec1 are the two with the highest penalties. This means the parametrization for the two ligand is not very successful and the error could lead to this outlying behavior. Actually, the simulated dissociation time is dependent on so many variables, such as crystallization quality, force field accuracy, statistical fluctuation, etc., it is probably not possible to get them all correctly. But the overall trend is clear and correct, which proves that KAMD is a suitable tool for study of inhibitory behaviors.

One more observation we can obtain from this test is that the KAMD time acceleration seems to follow a logarithmic rule. It seems that the KAMD dissociation time increases 100∼200ns each time the k_off_ rate drops one order of magnitude. We included a fitting plot between logarithmic reciprocal of k_off_ and the KAMD dissociation time, as shown in Figure S7. The fitting utilized the 7 successful values from the type-2 inhibitors. The fitting seems reasonable, and its R-square value is 0.669. Although we have not acquired any theory of KAMD time acceleration, a logarithmic rule seems a convincing model. Applying the fitting result to type-1 inhibitors and using 1qbs result as reference gives the extrapolated KAMD dissociation time for 1g2k/1g35 of about 1060 ns, agreeing with the simulation result. Of course, further tests and verifications are needed to confirm this.

## Conclusion

In this work, we combined the atomic accuracy of MD and diffusion behavior of BD to give a new type of enhanced sampling method, KAMD. KAMD utilizes re-oriented velocities in MD to accelerate domain/motif level of sampling. The special velocity distributions represent events that, though rare, do occur in standard MD simulations, making KAMD faithfully reproduce MD motions over large time scale. Also, it can be proved that KAMD preserves free energy surface and local time evolution.

In the Alanine dipeptide example, we showed the three key features of KAMD: 1. We showed that KAMD preserves the free energy surface by presenting the Ramachandran plot from both classic MD and KAMD simulations. 2. We showed that KAMD samples about 10-fold faster than MD 3. We showed that KAMD has similar time evolution behavior as MD, in a short period of time.

In the TSPI polypeptide and the thrombin test case, we showed that KAMD can realize the conformational change that happens at millisecond time scale in experiments. The trans-to-cis conformational change of TSPI and close-to-open conformational change of thrombin both can be simulated by KAMD, with 1us simulation. These tests proved that KAMD can accelerate the kinetics by thousands of folds.

In the test of Immunoglobulin E, we compared the PCA results from MD and KAMD simulation of free IgE-Fc system. The PCA showed a 5-time larger sampling area of KAMD. We also used the KAMD simulation of both free IgE-Fc and IgE-Fc-sFcεRIα complex systems to compare with fluorescence experiments. The results showed that KAMD simulation reproduces the experiments perfectly, while the conventional MD simulation cannot provide more information than the crystal structure.

Finally, in the HIV-1 protease inhibitor unbinding tests, we showed that KAMD can be applied for k_off_ rate studies. The KAMD relative dissociation time reproduced the k_off_ rate trend of the inhibitors. The simulated unbinding kinetics seems follow a logarithmic law and the fitting curve proved it.

In sum, we showed that KAMD is an excellent simulation tool for large-scale conformational changes and ligand unbinding events. The kinetics are accelerated thousands of folds and the accuracy is comparable to experiments. With full development, KAMD should prove widely useful for scientific discovery in molecular biology.

## Data and Software Availability Statement

The software and data that support the findings of this study are available from the corresponding author upon reasonable request. The software will also be available publicly in next AMBER release.

## Author Information Corresponding Author

Haixin Wei − Department of Chemistry and Biochemistry, University of California, San Diego, La Jolla, California 92093, United States;

## Acknowledgements

This work used Hopper at the Triton Shared Computing Cluster at the San Diego Supercomputer Center. The author thanks Prof. J. Andrew McCammon for helpful discussions and extensive revisions of the manuscript.

