## supporting information for "Enhanced Sampling on Domain/Motif Level with Kinetic Accelerated Molecular Dynamics"

### KAMD preserves conformational free energy

Assume canonical ensemble, the probability function of each microstate is:

$$e^{-\frac{E_{total}}{KT}} = e^{-\frac{U}{KT}} \cdot e^{-\frac{E}{KT}}$$

$U$ is the system total potential energy, and $E$ is the system total kinetic energy. $U$ is the same between a MD and a KAMD system, because we only altered velocities of marked atoms.

We only care about the probability distributions of macromolecules (biomolecules), thus:

$$P(state\ of\ macromolecule) = \int_{solvent} e^{-\frac{U}{KT}} \cdot \int_{solvent} e^{-\frac{E}{KT}} dxdp$$

$$= \int_{solvent} e^{-\frac{U}{KT}} \cdot e^{-\frac{E_{macromolecule}}{KT}} \cdot \int_{other\ solvent} e^{-\frac{E_{other\ solvent}}{KT}} \cdot \int_{marked-solvent} e^{-\frac{E_{marked-solvent}}{KT}} dxdp$$

Without loss of generality, we assume all the solvent molecules are interchangeable, though there are waters, ions, etc. Also, because the rigidity of input "regions", we assume that the marked solvents (and of course, the marked atoms in macromolecule) are of a consistent number each BD step.

Based on the two assumptions:

$$P(state\ of\ macromolecule)$$

$$\approx \int_{solvent} e^{-\frac{U}{KT}} \cdot e^{-\frac{E_{macromolecule}}{KT}} \cdot \int_{other\ solvent} e^{-\frac{E_{other\ solvent}}{KT}} \cdot \int_{1}^{n_{marked-solvent}} e^{-\frac{E_{marked-solvent}}{KT}} dxdp$$

$$= \int_{solvent} e^{-\frac{U}{KT}} \cdot e^{-\frac{E_{macromolecule}}{KT}} \cdot \int_{other\ solvent} e^{-\frac{E_{other\ solvent}}{KT}} \cdot \int_{1}^{n_{marked-solvent}} e^{-\frac{\sum\frac{1}{2}mv^2}{KT}} dxdp$$

In the last line, the first three factors are the same for normal MD systems and KAMD systems, and the difference comes from the last factor.

For the normal MD system:

$$\int_{1}^{n_{marked-solvent}} e^{-\frac{\sum\frac{1}{2}mv^2}{KT}} d\vec{v} = \prod \int e^{-\frac{\frac{1}{2}mv^2}{KT}} d\vec{v} = \prod \int e^{-\frac{\frac{1}{2}mv^2}{KT}} v^2 dv d\hat{n}$$

$$= (4\pi)^{n_{marked-solvent}} \prod \int e^{-\frac{\frac{1}{2}mv^2}{KT}} v^2 dv$$

$\hat{n}$ here is a unit vector.

For simplicity, we assume the KAMD system to have only one "region", so all the marked solvent velocities aligned with one random direction:

$$\int_1^{n_{marked-solvent}} e^{-\frac{\sum\frac{1}{2}mv^2}{KT}} d\vec{v}$$
$$= \int e^{-\frac{\frac{1}{2}m{v_1}^2}{KT}} {v_1}^2 dv_1 d\hat{n}_1 \cdot \prod_2^{n_{marked-solvent}} \int e^{-\frac{\frac{1}{2}mv^2}{KT}} \delta(\hat{n} - \hat{n}_1) v^2 dv d\hat{n}$$

Here, we used $\hat{n}_1$ as the chosen random direction $\hat{n}_{random}$, and $\delta$ is the Dirac delta function.

$$\int_1^{n_{marked-solvent}} e^{-\frac{\sum\frac{1}{2}mv^2}{KT}} d\vec{v} = (4\pi) \cdot \int e^{-\frac{\frac{1}{2}m{v_1}^2}{KT}} {v_1}^2 dv_1 \cdot \prod_2^{n_{marked-solvent}} \int e^{-\frac{\frac{1}{2}mv^2}{KT}} v^2 dv$$
$$= (4\pi) \prod \int e^{-\frac{\frac{1}{2}mv^2}{KT}} v^2 dv$$

Right now, we have, for MD system:

$$P(state\ of\ macromolecule)$$
$$= \int_{solvent} e^{-\frac{U}{KT}} \cdot e^{-\frac{E_{macromolecule}}{KT}} \cdot \int_{other\ solvent} e^{-\frac{E_{other\ solvent}}{KT}}$$
$$\cdot (4\pi)^{n_{marked-solvent}} \prod \int e^{-\frac{\frac{1}{2}mv^2}{KT}} v^2 dv$$

For KAMD system:

$$P(state\ of\ macromolecule)$$
$$= \int_{solvent} e^{-\frac{U}{KT}} \cdot e^{-\frac{E_{macromolecule}}{KT}} \cdot \int_{other\ solvent} e^{-\frac{E_{other\ solvent}}{KT}}$$
$$\cdot (4\pi) \prod \int e^{-\frac{\frac{1}{2}mv^2}{KT}} v^2 dv$$

Clearly, we can see that the macromolecule state probability function between normal MD systems and KAMD systems differ by only a constant factor $(4\pi)^{n_{marked-solvent}-1}$.

Next, we integral on velocities of the macromolecule to obtain the conformation probability function:

$$P(conformation\ of\ macromolecule) = \int_{velocities} P(state\ of\ macromolecule)$$

The only factor in $P(state\ of\ macromolecule)$ that related to this step is:

$$\int_{velocities} e^{-\frac{E_{macromolecule}}{KT}} d\vec{v}$$

Using a similar procedure, we can obtain, for MD system:

$$\int_{velocities} e^{-\frac{E_{macromolecule}}{KT}} d\vec{v} = \int_{macromolecule-not-marked} e^{-\frac{E_{macromolecule-not-marked}}{KT}} \cdot (4\pi)^{n_{marked-macromolecule}} \prod \int e^{-\frac{\frac{1}{2}mv^2}{KT}} v^2 dv$$

For KAMD system:

$$\int_{velocities} e^{-\frac{E_{macromolecule}}{KT}} d\vec{v} = \int_{macromolecule-not-marked} e^{-\frac{E_{macromolecule-not-marked}}{KT}} \cdot \prod \int e^{-\frac{\frac{1}{2}mv^2}{KT}} v^2 dv$$

Noted that, there is no $(4\pi)$ factor in KAMD expression because the velocity direction is determined by the marked solvent part.

Taking all together, we reach the conclusion:

$$P(MD\ conformation\ of\ macromolecule) \approx (4\pi)^{n_{marked}-1} P(KAMD\ conformation\ of\ macromolecule)$$

The conformational distribution of MD and KAMD differ by only a constant $(4\pi)^{n_{marked}-1}$. Thus, the conformational free energy of MD and KAMD is the same.

**KAMD preserves time evolution approximately**

To simplify the analysis, we assume there is only one input region. Since each region will have collective motions, like rigid bodies, we assume the state of each input region can be approximately represented by six variable and their momentum:

$$(x, y, z, \varphi, \phi, \tau, p_x, p_y, p_z, p_\varphi, p_\phi, p_\tau)$$

Since KAMD would not affect rotational motion, the angel part can be integrated and ignored. Thus, for MD systems, the conformational distribution function of an input region can be written as:

$$P(x, y, z, p_x, p_y, p_z) = e^{-\frac{U_{x,y,z}}{KT}} \cdot e^{-\frac{\frac{1}{2}\frac{p^2}{m_{eff}}}{KT}}$$

$U_{x,y,z}$ is the averaged potential energy, and $m_{eff}$ is the effective mass the input region.

However, the distribution will be altered by KAMD, since the marked velocities are aligned to one direction. For KAMD systems, the probability distribution function should be:

$$P(x, y, z, p_x, p_y, p_z) = e^{-\frac{U_{x,y,z}}{KT}} \cdot (c_1 e^{-\frac{\frac{1}{2}\frac{p^2}{m'_{eff}}}{KT}} + c_2 D(p))$$

Here, the velocity distribution is sum of two parts: the marked atoms and the unmarked atoms, and $c_1$ and $c_2$ are constant factors representing proportions of the two parts. Since KAMD will not alter velocities of unmarked atoms, their velocity distribution is similar to MD systems. For the marked atoms, their velocity distribution will be a complicated function. For example, the sum of two aligned Gaussian distribution will follow the distribution:

$$D\big(x|_{\,summation\ of\ 2}\big) = e^{-\frac{1}{2}x^2} \cdot \text{erf}\,(\frac{|x|}{\sqrt{2}})$$

Although the exact formulation cannot be obtained, $D(p)$ shows two obvious properties:
1. $D(p)$ is an even function, for all three variables
2. unlike normal Gaussian distribution, the highest probability value does not happen at $p = 0$. On the contrary, $D(p = 0) = 0$. This is easily seen from summation of 2 aligned Gaussian above.

Now, we will try to prove the time evolution of the system is preserved with KAMD, but only in a local condition.

Suppose at beginning, the input region is at conformation $(x, y, z)$. Thus, for a MD system, the starting distribution at phase space should be:

$$P(MD) = \delta(x, y, z) \cdot e^{-\frac{\frac{1}{2}\frac{p^2}{m_{eff}}}{KT}}$$

And that of KAMD system should be:

$$P(MD) = \delta(x, y, z) \cdot (c_1 e^{-\frac{\frac{1}{2}\frac{p^2}{m'_{eff}}}{KT}} + c_2 D(p))$$

Given time, the two distributions will evolve in the phase space. Off course, after evolution, the end time distribution is different for them. However, we will show that, the center (average) of the two distributions, will be the same, if consider a short period of time.

The center (average) of the distribution:

For MD system: $\int \vec{q}\,\delta(x, y, z) \cdot e^{-\frac{\frac{1}{2}\frac{p^2}{m_{eff}}}{KT}} dqdp = (x, y, z) = \vec{q}_{t=0}$

For KAMD system: $\int \vec{q}\,\delta(x, y, z) \cdot (c_1 e^{-\frac{\frac{1}{2}\frac{p^2}{m'_{eff}}}{KT}} + c_2 D(p)) dqdp = (x, y, z) = \vec{q}_{t=0}$

Compute their first order derivative:

For MD system: $\frac{d}{dt}\int \vec{q}\,\delta(x, y, z) \cdot e^{-\frac{\frac{1}{2}\frac{p^2}{m_{eff}}}{KT}} dqdp = \int \frac{d\vec{q}}{dt}\delta(x, y, z) \cdot e^{-\frac{\frac{1}{2}\frac{p^2}{m_{eff}}}{KT}} dqdp =$
$\int \frac{\vec{p}}{m}\delta(x, y, z) \cdot e^{-\frac{\frac{1}{2}\frac{p^2}{m_{eff}}}{KT}} dqdp = 0$

For KAMD system: $\frac{d}{dt}\int \vec{q}\,\delta(x,y,z)\cdot(c_1 e^{-\frac{\frac{1}{2}\frac{p^2}{m'_{eff}}}{KT}} + c_2 D(p))dqdp = \int \frac{d\vec{q}}{dt}\delta(x,y,z)\cdot (c_1 e^{-\frac{\frac{1}{2}\frac{p^2}{m'_{eff}}}{KT}} + c_2 D(p))dqdp = \int \frac{\vec{p}}{m}\delta(x,y,z)\cdot(c_1 e^{-\frac{\frac{1}{2}\frac{p^2}{m'_{eff}}}{KT}} + c_2 D(p))dqdp = 0$

In the calculation, the first step used Liouville's theorem, and the second step is because it is an integral of an odd function times an even function.

The second order derivative:

For MD system: $\frac{d^2}{dt^2}\int \vec{q}\,\delta(x,y,z)\cdot e^{-\frac{\frac{1}{2}\frac{p^2}{m_{eff}}}{KT}}dqdp = \int \frac{d\vec{p}}{mdt}\delta(x,y,z)\cdot e^{-\frac{\frac{1}{2}\frac{p^2}{m_{eff}}}{KT}}dqdp = \int \frac{\vec{F}}{m}\delta(x,y,z)\cdot e^{-\frac{\frac{1}{2}\frac{p^2}{m_{eff}}}{KT}}dqdp = \frac{\vec{F}(x,y,z)}{m}$

For KAMD system: $\frac{d^2}{dt^2}\int \vec{q}\,\delta(x,y,z)\cdot(c_1 e^{-\frac{\frac{1}{2}\frac{p^2}{m'_{eff}}}{KT}} + c_2 D(p))dqdp = \int \frac{d\vec{p}}{mdt}\delta(x,y,z)\cdot (c_1 e^{-\frac{\frac{1}{2}\frac{p^2}{m'_{eff}}}{KT}} + c_2 D(p))dqdp = \int \frac{\vec{F}}{m}\delta(x,y,z)\cdot(c_1 e^{-\frac{\frac{1}{2}\frac{p^2}{m'_{eff}}}{KT}} + c_2 D(p))dqdp = \frac{\vec{F}(x,y,z)}{m}$

Here, $\vec{F}(x,y,z)$ is the total force at the starting structure. Again, the computation utilized Liouville's theorem, twice actually.

The third order derivative:

For MD system: $\frac{d^3}{dt^3}\int \vec{q}\,\delta(x,y,z)\cdot e^{-\frac{\frac{1}{2}\frac{p^2}{m_{eff}}}{KT}}dqdp = \int \frac{d\vec{F}}{mdt}\delta(x,y,z)\cdot e^{-\frac{\frac{1}{2}\frac{p^2}{m_{eff}}}{KT}}dqdp = \int \frac{(\nabla\vec{F})\cdot\vec{p}}{m^2dt}\delta(x,y,z)\cdot e^{-\frac{\frac{1}{2}\frac{p^2}{m_{eff}}}{KT}}dqdp = 0$

For KAMD system: $\frac{d^3}{dt^3}\int \vec{q}\,\delta(x,y,z)\cdot(c_1 e^{-\frac{\frac{1}{2}\frac{p^2}{m'_{eff}}}{KT}} + c_2 D(p))dqdp = \int \frac{d\vec{F}}{mdt}\delta(x,y,z)\cdot (c_1 e^{-\frac{\frac{1}{2}\frac{p^2}{m'_{eff}}}{KT}} + c_2 D(p))dqdp = \int \frac{(\nabla\vec{F})\cdot\vec{p}}{m^2dt}\delta(x,y,z)\cdot(c_1 e^{-\frac{\frac{1}{2}\frac{p^2}{m'_{eff}}}{KT}} + c_2 D(p))dqdp = 0$

Starting from the fourth order, the derivatives no longer are the same for the two systems.

From above calculation, we can see that, the center of the time evolved distribution of MD and KAMD differ only from the fourth order. Thus, in a short period of time, or a local conformational space, the time evolution path is preserved with KAMD.

**Simulation details**

1. Force field: The alanine dipeptide and TSPI polypeptide systems were built in AMBER tleap program and parametrized with force field, ff19SB. The thrombin, Immunoglobulin E, and HIV-1 protease systems were built with CHARMM-GUI and parametrized with CHARMM36m force field. The ligands in HIV-1 protease systems were parametrized with CHARMM general force field. The protonation state of the ligands was determined with Schrodinger Maestro software. The lipid bilayer in the IgE-Fc-sFcεRIα complex was built by CHARMM-GUI, with a composition of 67% POPC and 33% cholesterol.

2. Solation: All the systems were solvated with TIP3P waters and 0.15 M NaCl. 10 Å space was added to the edge of each system, except for the free IgE-Fc system, in which 20 Å space was added.

3. Minimization, heating, and equilibration: For all the system, 100000 steps minimization was first done by AMBER PMEMD.CUDA_DPFP, then, another 100000 steps minimization was done with AMBER PMEMD.CUDA_SPFP. After minimization, each system was gradually heated from 10 to 300 K over 1 ns in NVT condition, except TSPI and HIV-1 protease systems. For TSPI system, it was gradually heated from 10 to 400k over 2ns, and for HIV-1 protease systems, they were gradually heated from 10 to 293.15k over 1ns. After heating, each system was equilibrated under 1bar and the temperature from heating, for another 1 ns, in an NPT ensemble. The NPT condition was controlled using a Langevin thermostat with a friction coefficient γ of 5.0 $ps^{-1}$ and a Berendsen barostat with a time constant τ of 2.0 ps.

4. Production KAMD simulation: 10 independent KAMD trajectories were run for each system, restarting from the previous equilibration. All the KAMD simulations were done under NVT condition, using a Langevin thermostat with a friction coefficient γ of 5.0 $ps^{-1}$. For Alanine dipeptide and Immunoglobulin E systems, each trajectory was run for 200ns. For TSPI system, each trajectory was run for 2us. For thrombin and HIV-1 protease systems, each trajectory was run for 1us.

**Supplementary figures**

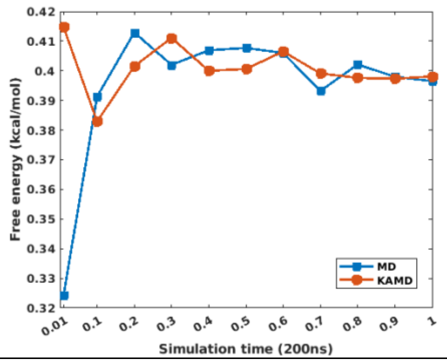


(a)

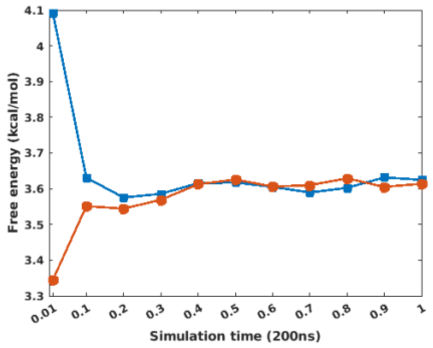


(b)

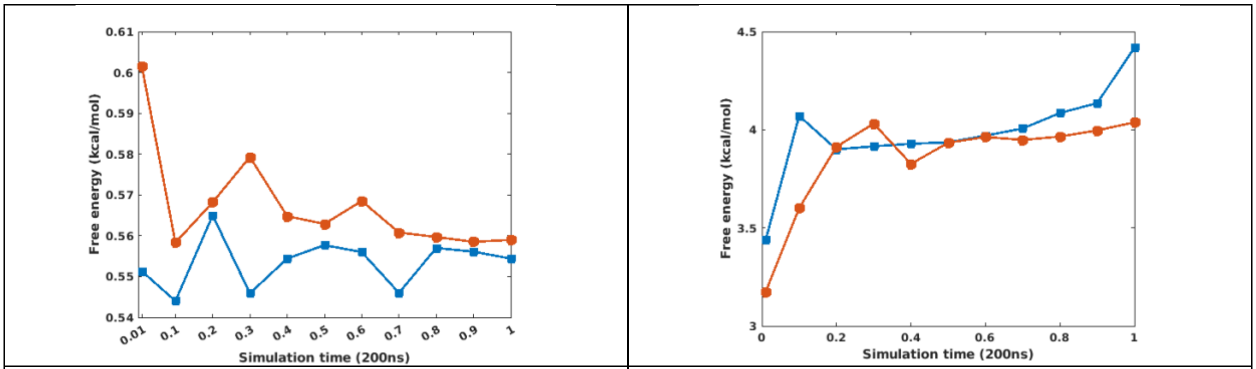


(c) (d)

Figure S1. Free energy convergence plot for other conformations. (a) $\alpha$ conformation: Phi = -75° and Psi = 20°. (b) $\beta$ conformation: Phi = -150° and Psi = -130°. (c) Phi = -60° and Psi = -40°. (d) Phi = 70° and Psi = -50°.

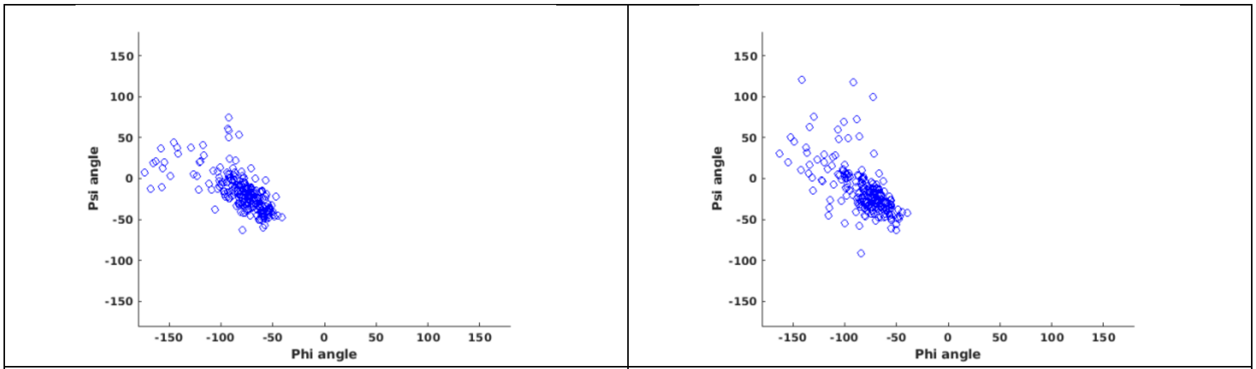


(a) (b)

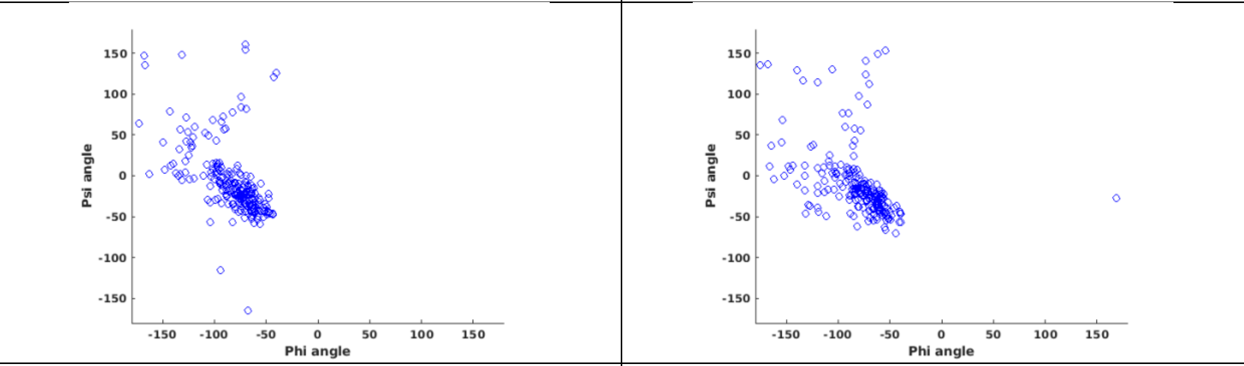


(c) (d)

| | |
|---|---|
| 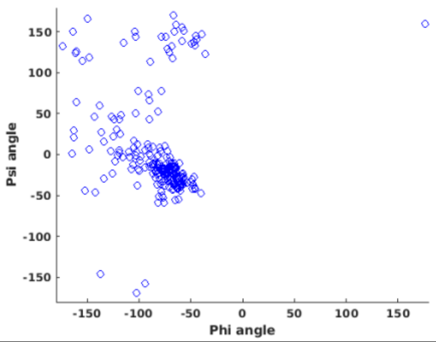 | 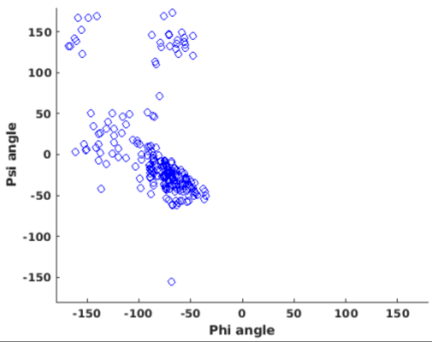 |
| (e) | (f) |



Figure S2. Conformation distribution after MD/KAMD simulation, from $\varepsilon$ conformation. (a) 4ps MD simulation (b) 4ps KAMD simulation (c) 10ps MD simulation (d) 10ps KAMD simulation (e) 20ps MD simulation (f) 20ps KAMD simulation

| | |
|---|---|
| 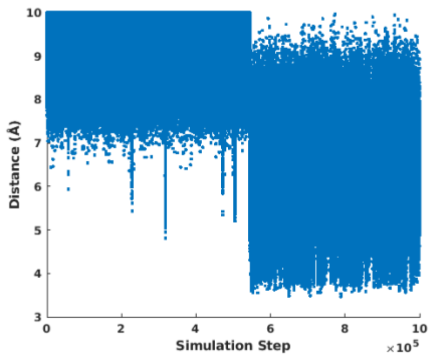 | 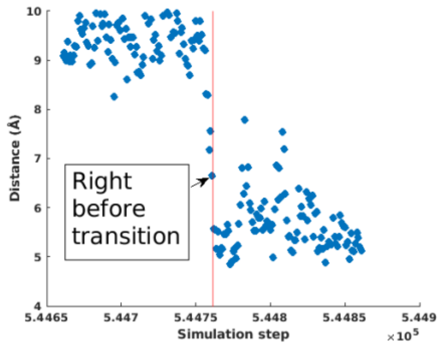 |
| (a) | (b) |



Figure S3. Distance between Threonine and Isoleucine C-$\alpha$ atoms, plot against simulation steps. (a) The distance distribution of the entire trajectory. (b) The distance distribution of 100ps before and after the cis-trans conformational change.

| | |
|---|---|
| 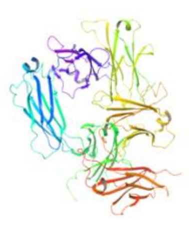 | 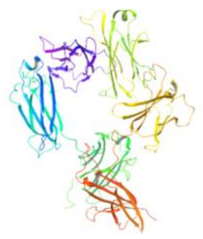 |
| (a) | (b) |

Figure S4. Representative structure from trajectory clustering. (a) Representative structure of MD trajectories. (b) Representative structure of KAMD trajectories.

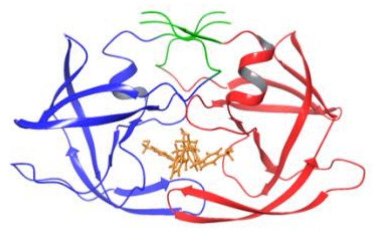

(a)

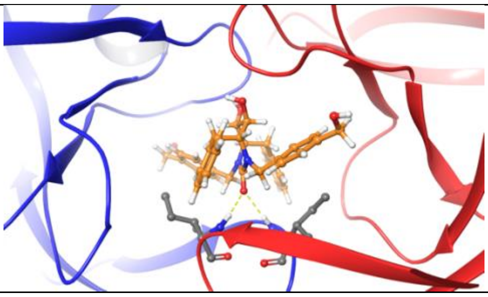

(b)

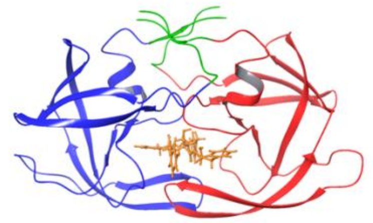

(c)

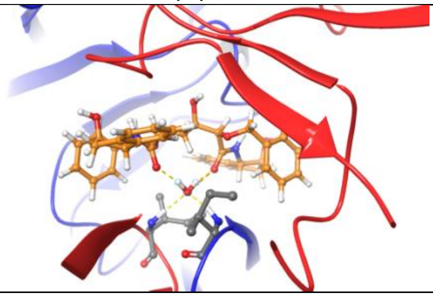

(d)

Figure S5. Input “regions” and active site view of HIV-1 protease inhibitors. (a) Input “regions” of type-1 inhibitor systems. The red, blue, and yellow part are input “regions”. The green part is not in any input “regions”. (b) Type-1 inhibitor active site. (c) Input “regions” of type-2 inhibitor systems. The red, blue, and yellow part are input “regions”. The green part is not in any input “regions”. (d) Type-2 inhibitor active site.

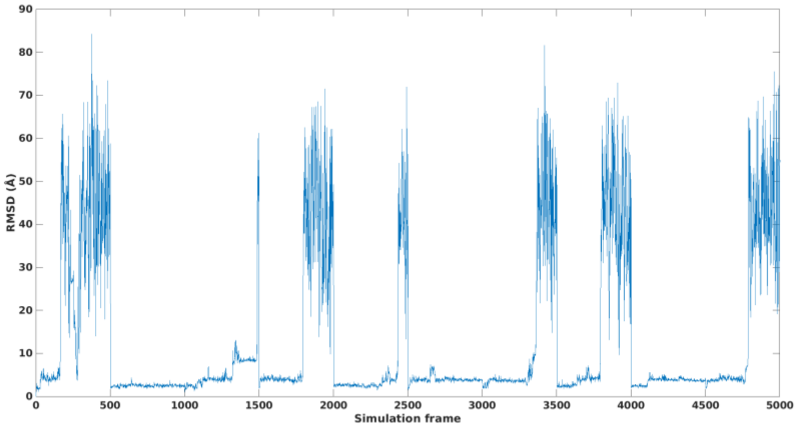


(a)

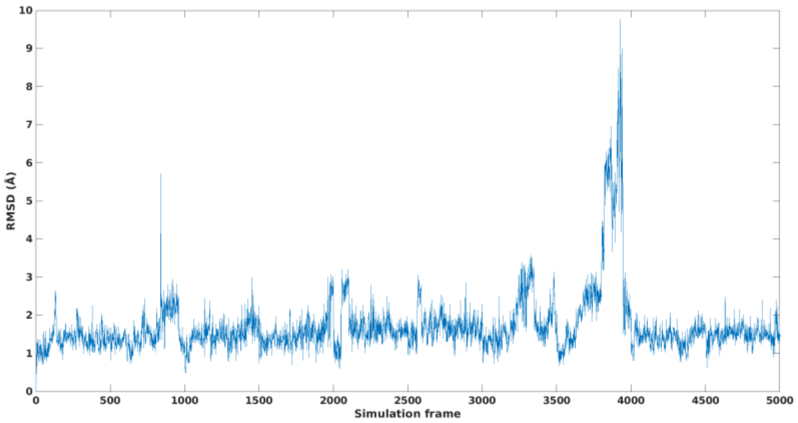
RMSD (Å)
Simulation frame
0
500
1000
1500
2000
2500
3000
3500
4000
4500
5000
0
1
2
3
4
5
6
7
8
9
10

(b)

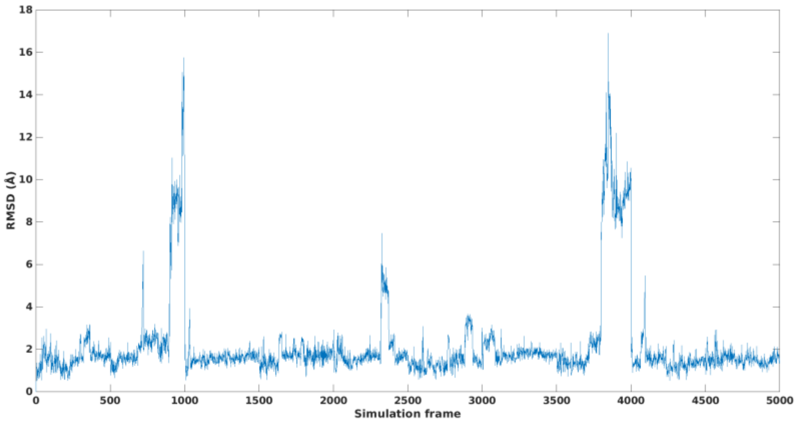
RMSD (Å)
Simulation frame
0
500
1000
1500
2000
2500
3000
3500
4000
4500
5000
0
2
4
6
8
10
12
14
16
18

(c)

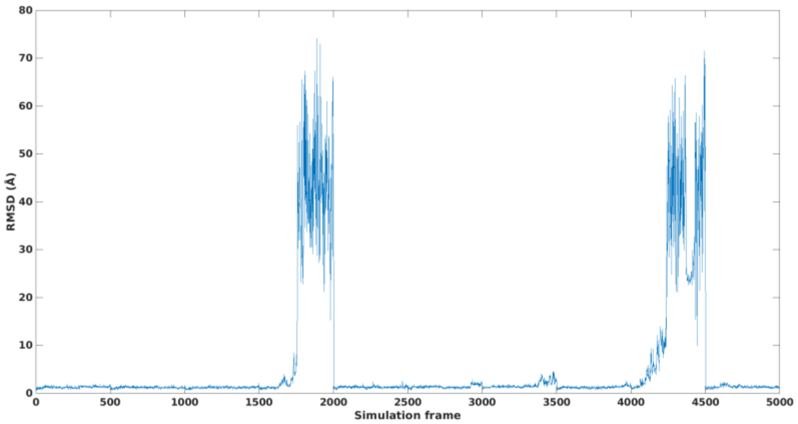
RMSD (Å)
Simulation frame
0
500
1000
1500
2000
2500
3000
3500
4000
4500
5000
0
10
20
30
40
50
60
70
80

(d)

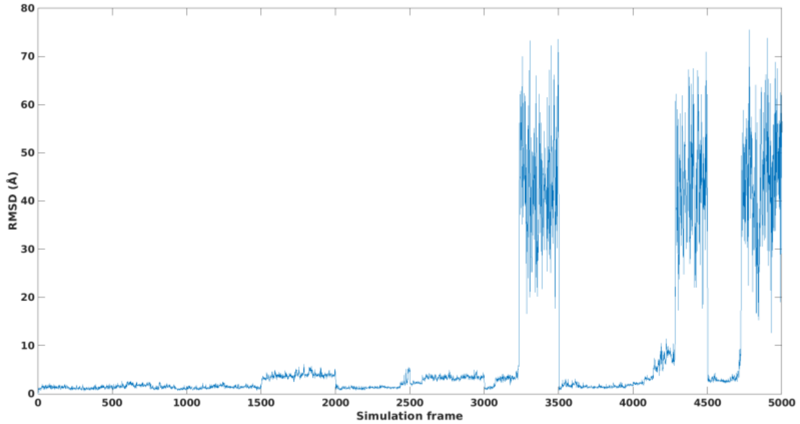
RMSD (Å)
Simulation frame
0
10
20
30
40
50
60
70
80
500
1000
1500
2000
2500
3000
3500
4000
4500
5000

(e)

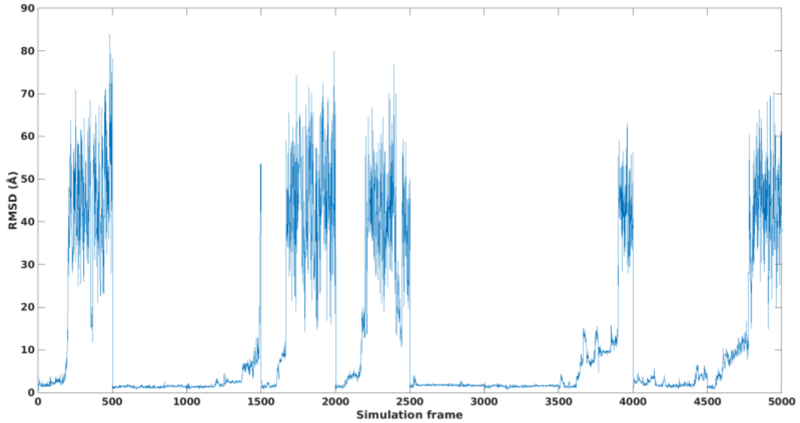
RMSD (Å)
Simulation frame
0
10
20
30
40
50
60
70
80
90
500
1000
1500
2000
2500
3000
3500
4000
4500
5000

(f)

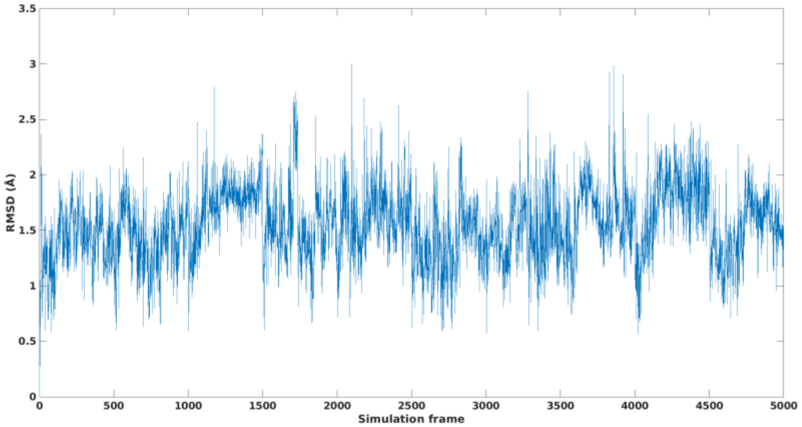
RMSD (Å)
Simulation frame
0
0.5
1
1.5
2
2.5
3
3.5
500
1000
1500
2000
2500
3000
3500
4000
4500
5000

(g)

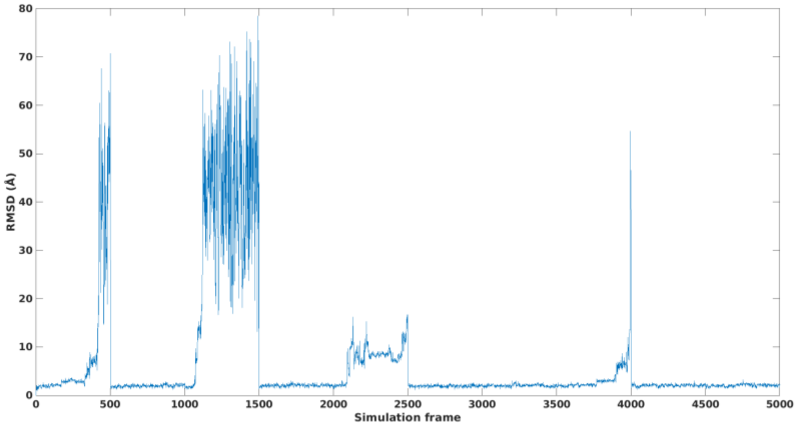

RMSD (Å)
Simulation frame


(h)

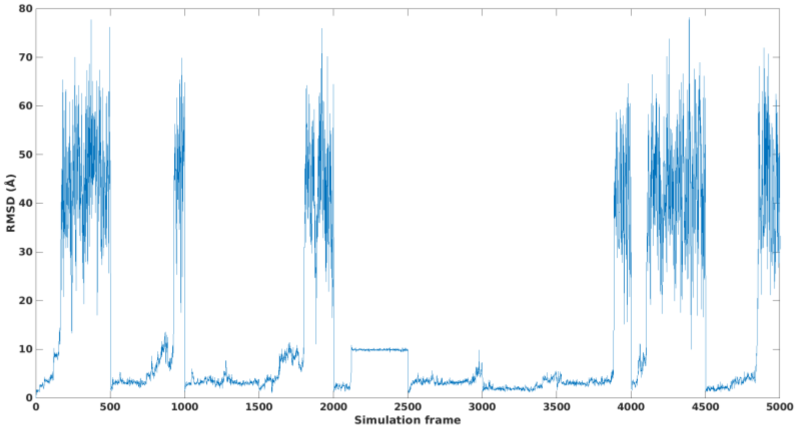

RMSD (Å)
Simulation frame


(i)

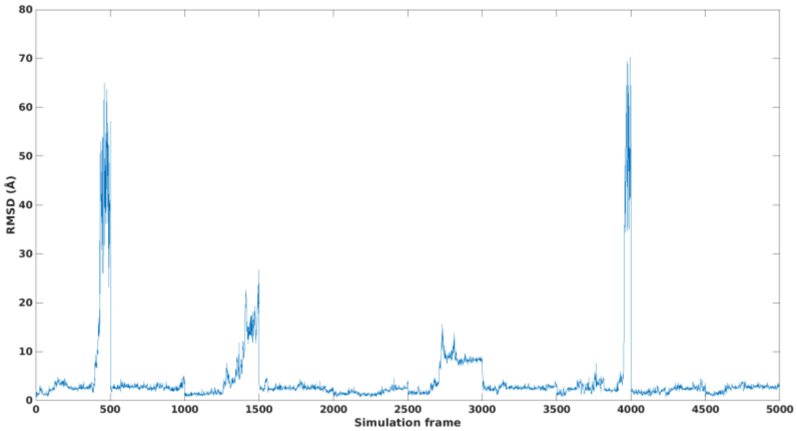

RMSD (Å)
Simulation frame


(j)

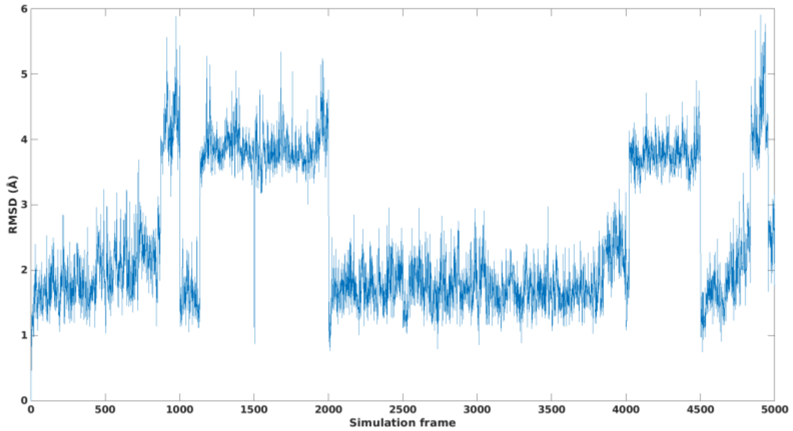


(k)

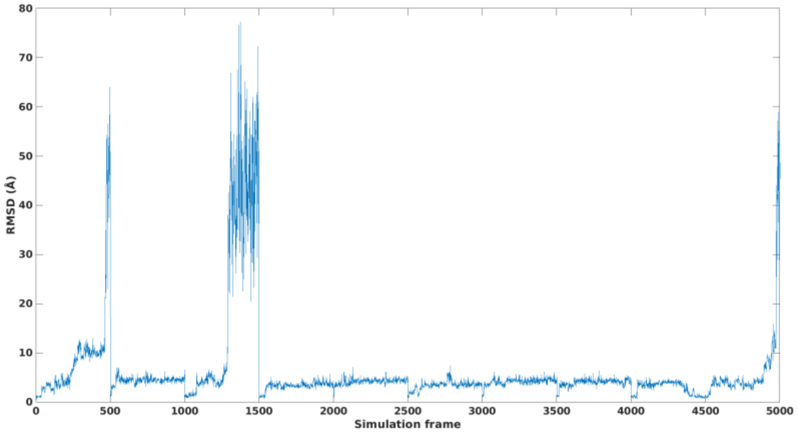


(l)

Figure S6. Ligand RMSD plot of all 12 HIV-1 protease inhibitor systems. (a) 1qbs (b) 1g2k (c) 1g35 (d) 1d4i (e) 1eby (f) 1d4h (g) 3ekv (h) 1hxw (i) 2bpx (j) 1ohr (k) 1ec1 (l) 3oxc

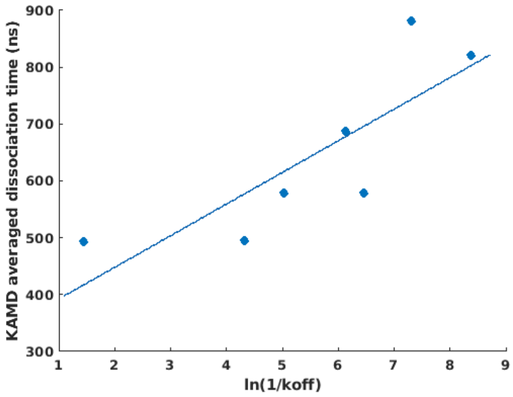


Figure S7. Linear fitting between ln(1/ $K_{off}$) and KAMD dissociation time.

**Supplementary tables**

| Trajectory number | IgE-Fc-sFcεRIα complex simulation frame / major structural event |
|---|---|
| 1 | 700 / complex dissociation |
| 2 | 500 / complex dissociation |
| 3 | None |
| 4 | 900 / Cε4 dimer uncoupling |
| 5 | None |
| 6 | 300 / complex dissociation |
| 7 | None |
| 8 | None |
| 9 | None |
| 10 | None |

Table S1. Major structural event and the simulation frame it happened in the IgE-Fc-sFcεRIα complex system. For both free IgE-Fc system and IgE-Fc-sFcεRIα complex system, 200ns KAMD simulation was done, and 1000 frames was written out.

| PDB ID | Parameter penalty | Charge penalty |
|---|---|---|
| 1qbs | 30.6 | 16.0 |
| 1g2k | 20.8 | 11.8 |
| 1g35 | 32.5 | 17.6 |
| 1d4i | 72.5 | 26.3 |
| 1eby | 72.5 | 26.3 |
| 1d4h | 72.5 | 26.3 |
| 3ekv | 76.6 | 11.1 |
| 1hxw | 48.5 | 35.0 |
| 2bpx | 72.5 | 35.7 |
| 1ohr | 63 | 25.0 |
| 1ec1 | 81 | 37.4 |
| 3oxc | 37.9 | 30.1 |

Table S2. CHARMM ligand parametrization penalty of 12 HIV-1 protease inhibitor systems.
